# Disentangling locus of perceptual learning in the visual hierarchy of motion processing

**DOI:** 10.1101/282319

**Authors:** Ruyuan Zhang, Duje Tadin

## Abstract

Visual perceptual learning (VPL) can lead to long-lasting perceptual improvements. While the efficacy of VPL is well established, there is still a considerable debate about what mechanisms underlie the effects of VPL. Much of this debate concentrates on where along the visual processing hierarchy behaviorally relevant plasticity takes place. Here, we aimed to tackle this question in context of motion processing, a domain where links between behavior and processing hierarchy are well established. Specifically, we took advantage of an established transition from component-dependent representations at the earliest level to pattern-dependent representations at the middle-level of cortical motion processing. We trained two groups of participants on the same motion direction identification task using either grating or plaid stimuli. A set of pre- and post-training tests was used to determine the degree of learning specificity and generalizability. This approach allowed us to disentangle contributions from both low- and mid-level motion processing, as well as high-level cognitive changes. We observed a complete bi-directional transfer of learning between component and pattern stimuli as long as they shared the same apparent motion direction. This result indicates learning-induced plasticity at intermediate levels of motion processing. Moreover, we found that motion VPL is specific to the trained stimulus direction, speed, size, and contrast, highlighting the pivotal role of basic visual features in VPL, and diminishing the possibility of non-sensory decision-level enhancements. Taken together, our study psychophysically examined a variety of factors mediating motion VPL, and demonstrated that motion VPL most likely alters visual computation in the middle stage of motion processing.

## INTRODUCTION

A large body of evidence has shown that the human visual system can gain long-lasting perceptual improvements following several sessions of perceptual training. This phenomenon, called visual perceptual learning (VPL), has been an active area of research because VPL is a remarkable demonstration that human vision can remain plastic even in adulthood ^1,2^. Numerous studies have revealed training-induced perceptual improvements on a wide range of visual tasks, including low-level contrast and orientation discrimination tasks ^3,4,5,6^, mid-level motion and form tasks ^7,8,9^ and even high-level object and face recognition tasks ^10,11^.

While the robustness of learning effects is well established, debate persists with respect to the mechanisms underlying VPL. Early psychophysical work found that learning effects are usually confined to the trained parameters ^6,12^. Such strong specificity suggests that VPL most likely takes place within low-level visual areas (e.g., V1 or V2) since neurons therein exhibit narrow ranges of spatial and feature selectivity (e.g., orientation, motion direction). Recent evidence, however, challenges this idea by revealing an increasing number of cases where the transfer of VPL is viable to novel stimulus conditions and tasks ^13,14^. This is consistent with an involvement of higher-level visual areas, wherein neurons usually respond to larger spatial areas and more complex stimulus features. Some studies even suggest the contributions from the brain areas that process non-sensory attributes. For instance, perceptual learning might manifest as the change of decision variables encoded in the prefrontal cortex ^15^. Alternatively, perceptual learning might facilitate encoding of abstract concepts representing basic visual features (e.g., orientation and contrast) ^16^ or lead to a better set of task-specific rules ^17^. Given that these theories postulate changes beyond canonical sensory mechanisms, we refer to them as ‘non-sensory’ learning processes.

The task of linking VPL to specific brain areas is complicated by the complex functional specializations of the brain. The brain includes multiple brain regions that are organized into a coarse, but richly interconnected hierarchy ^18,19^. Even a simple perceptual choice likely arises from the interplay among multiple brain regions. One strategy is to take advantage of visual processes where links between behavior and neural structures are well established. Here, we focus on VPL in context of motion perception, a perceptual domain where we have a relatively advanced understanding of different processing stages ^20^. In primates, neurons selective to motion direction first occur in the earliest cortical areas V1 and V2 ^21^. However, conscious motion perception is most closely linked to intermediate visual areas, such as MT and V3A. These areas contain a large portion of neurons showing strong preferential responses to different motion directions ^22,23,24,25^. In addition, perceptual decisions based on motion stimuli have been linked to several higher-level brain areas (e.g., lateral intraparietal cortex (LIP) and prefrontal cortex). These areas are often ascribed as “evidence accumulators” that integrate sensory information provided by the upstream motion processing units in order to form perceptual decisions and guide visual behaviors ^26,27^ (but see ref. ^28^). Finally, nonsensory attributes, such as task rules and decision strategies, encoded in high-level cognitive areas, can also mediate performance in motion perception tasks ^29^. This complex hierarchy can be operationalized as a symbolic three-layer network (Figure 1). This network consists of a low-level (e.g., V1/V2), a middle level (e.g., MT/V3A) and a high-level (e.g., LIP, prefrontal cortex) processing stage.

**Figure 1.**
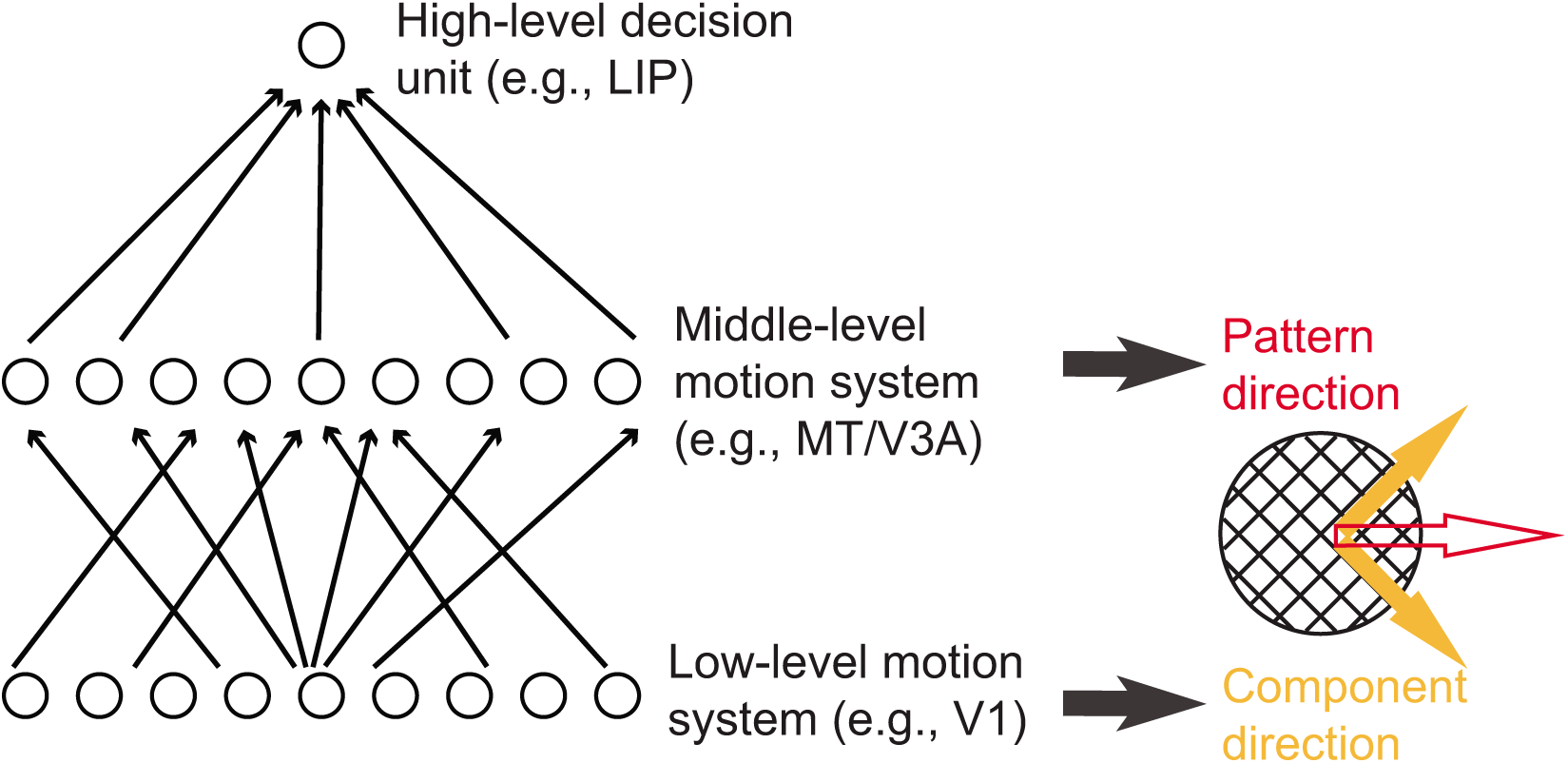
A simplified hierarchy of visual motion processing with three hierarchical stages ^20^. Neurons in the low-level motion system respond best to component directions of plaid stimuli (full orange arrows). Neurons in the middle-level motion system respond selectively to the perceived pattern motion direction (empty red arrow). In this and all subsequent figures, empty arrows indicate faster perceived speed associated with plaid motion. The third stage involves complex sensory and non-sensory high-level cognitive processes.

In contrast with the established understanding of visual motion processing stages, their role in human VPL is largely unknown. To address this question, we took advantage of an established transition from component-dependent representations at the earliest level to pattern-dependent representations in the middle-level of cortical motion processing ^30,31^. A plaid stimulus composed of two obliquely moving gratings (Figure 1) is generally perceived as a rigid object moving horizontally ^32^. While many MT neurons faithfully respond to the perceived motion direction in moving plaids, neurons in V1 primarily respond to the directions of two component gratings ^30,31^. This dissociation allows us to psychophysically infer the main locus of the behaviorally related plasticity induced by motion VPL. If learning effects are specific to the trained component motion, irrespective of the perceived pattern motion, it would indicate component-dependent learning predominantly at the lowest levels of motion processing. Conversely, if learning effects are specific to the perceived pattern motion, it would point toward pattern-dependent learning at middle-levels of motion processing. If this learning still retains some sensory-level specificity (e.g., speed, size and velocity), we can conclude that nonsensory cognitive processes are not the major drivers of the observed behavioral improvement.

## METHODS AND EXPERIMENTAL PROCEDURES

### Participants and apparatus

Fourteen undergraduate students from University of Rochester (18 to 22 years old, 5 males and 9 females) took part in this study. All participants had normal or corrected-to-normal vision. The Research Subjects Review Board at the University of Rochester approved experimental protocols and all participants provided written consent forms. Stimuli were generated by Matlab Psychtoolbox ^33^ and presented using customized digital light processing (DLP) projector (DepthQ WXGA 360 driven by a NVIDIA Quadro FX 4800 at 1280 × 720 resolution). The projector frame rate was 360 Hz, resulting in discrete 2.78-ms frames. DLP projectors are natively linear, and this was verified with a Minolta LS-110 photometer. Viewing distance was 61.5 inches, with a projected image of 46.74 × 25 inches.

### Stimulus and task settings

Participants were randomly assigned into two groups – one group trained on component motion (grating; N = 8) and another group trained on pattern motion stimuli (plaid; N = 6). All participants were tested and trained on a two-alternative forced choice motion direction identification task (Figure 2), reporting the perceived stimulus motion direction via key press. Auditory feedback was provided after each trial during the training phase but not at pre-/post-test (to minimize learning effects in pre-/post-test). To facilitate fixation, we used the following fixation sequence (Figure 2): a fixation circle (0.8° radius) appeared after each key press response and, the circle shrank to 0.13° over 200 ms, remained at that size for 360 ms, and then disappeared 360 ms before stimulus onset. We found in our previous work that this dynamic fixation sequence was very effective in guiding eye gaze to the center of the screen before the stimulus onset ^34^. The inter-trial interval was 1000 ms.

**Figure 2.**
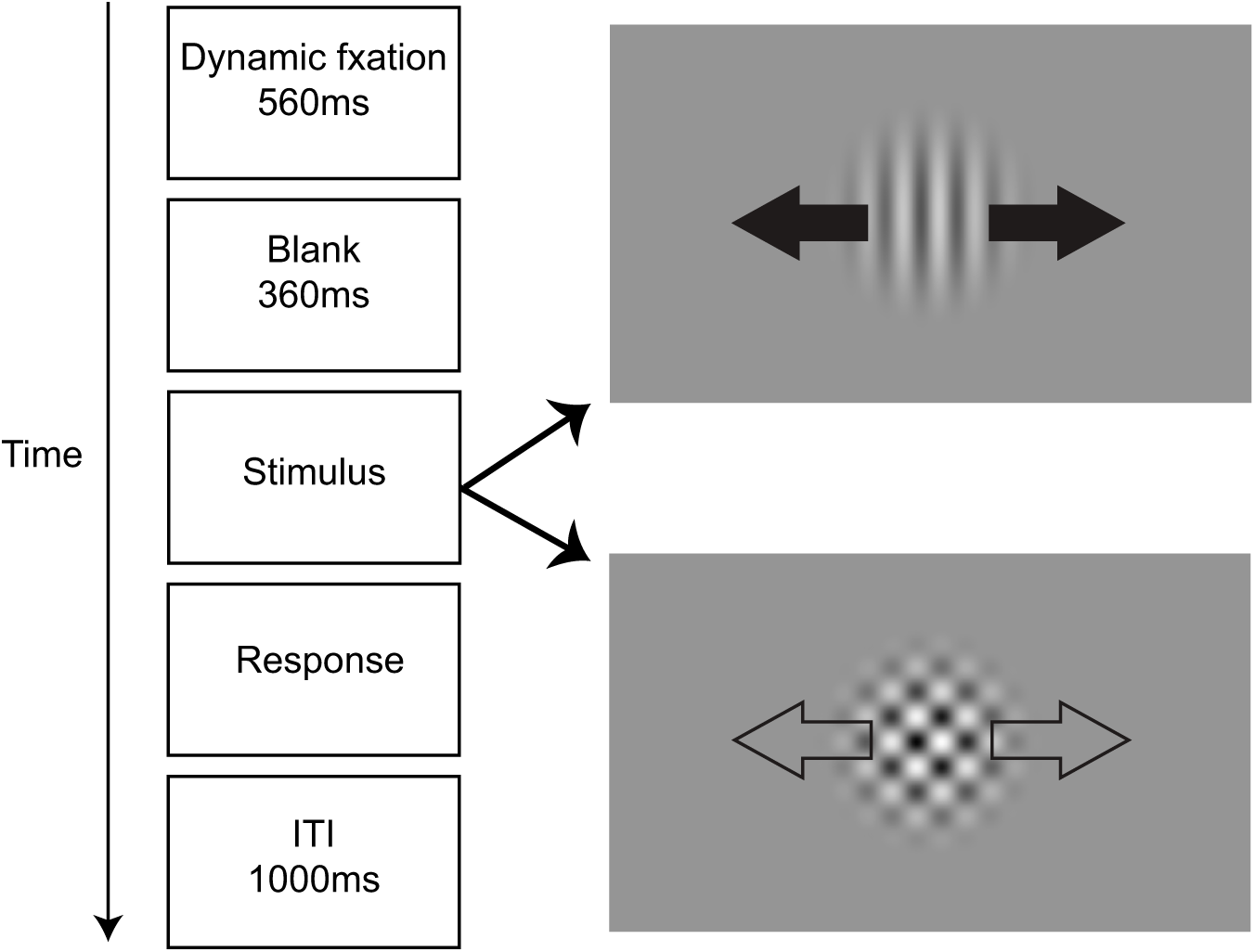
Task illustration showing trial structure used for all training, pre- and post-test conditions. Participants viewed a moving stimulus that was either a grating or a plaid (arrows are for illustration purposes only). Stimulus duration varied on each trial, as determined by two interleaved staircases. Participants indicated the perceived stimulus direction via button press (e.g., left vs. right in this case).

As detailed below, the two training groups used partially overlapping sets of pre- and post-test conditions. We selected this design to limit pre- and post-test sessions to only the most diagnostic test conditions for each group. This allowed us to test the bidirectional transfer between component and pattern motion, as well as the dependency of learning transfer on several key low-level stimulus features.

In the component-training group, the training stimulus was a grating (contrast = 50%, drift speed = 4°/s, radius = 8°, 2D raised cosine spatial envelope; spatial frequency = 1 cycle/°; Figure 3Aa). Training motion directions were either left/right or up/down, counterbalanced across participants. Motion directions for other stimulus conditions were adjusted according to the directions of trained stimuli. During the pre- and post-test, temporal duration thresholds (defined by the full-width at half-height of a hybrid between a Gaussian and a trapezoidal temporal envelope; see ^35^ for details) were measured across another five stimulus conditions: (1) a plaid stimulus moving in the trained directions (Figure 3Ab); (2) a plaid stimulus containing the trained component and moving 45° away from the trained direction (Figure 3Ac); (3-5) moving gratings that matched the trained grating except that they differed in (3) direction and orientation (orthogonal to the trained direction; Figure 3Ad), (4) stimulus size (radius = 1°, Figure 3Ae), and (5) contrast (contrast = 2%, Figure 3Af).

**Figure 3.**
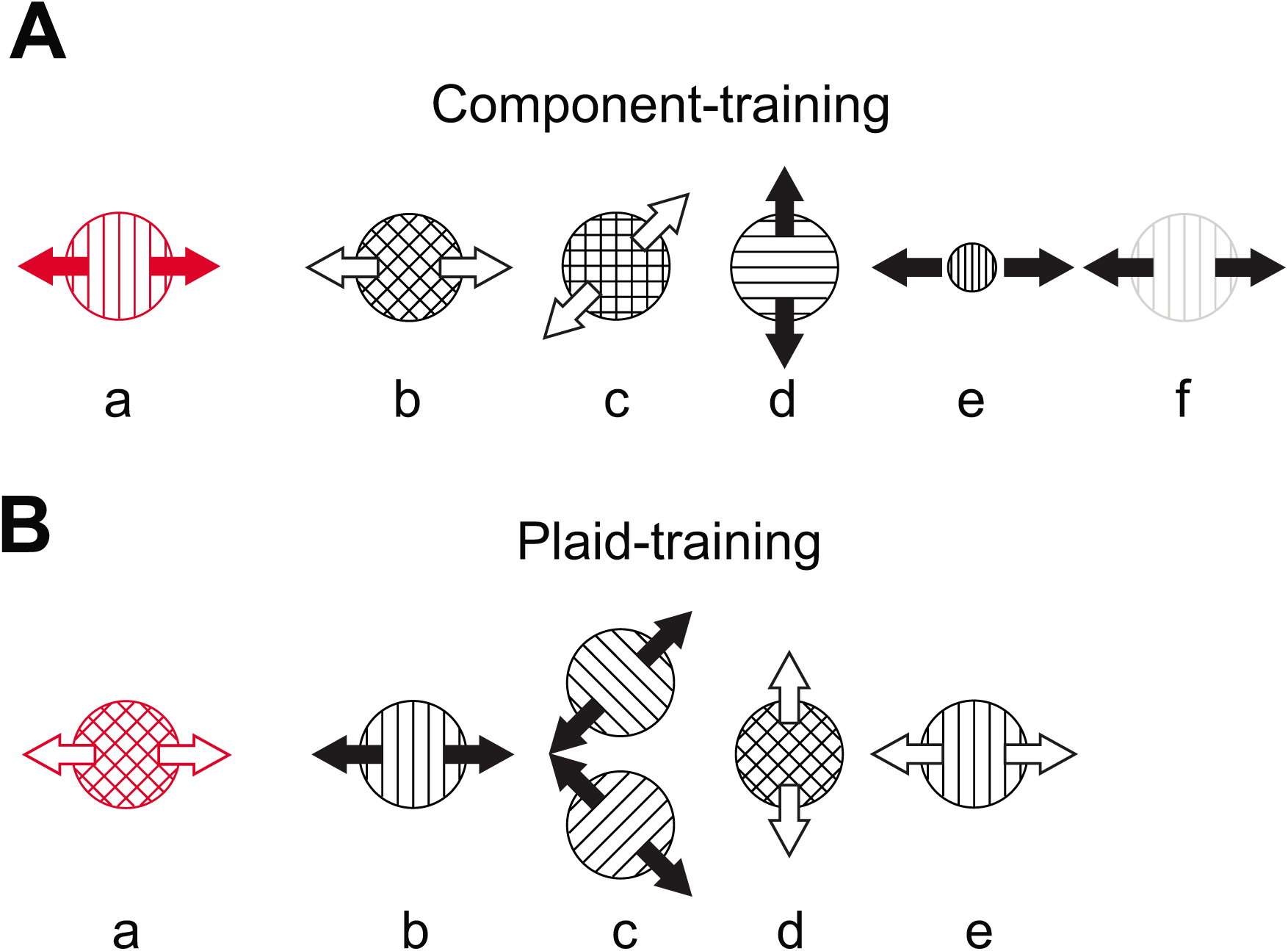
Pre- and post-training stimuli for (A) the component-training group and (B) the plaid-training group. The red icons show the training stimulus for each group. These conventions are kept in subsequent figures. With the exception of Be, the speed of all grating stimuli was 4°/s (marked by solid arrows). The plaid component speed was also 4°/s, which resulted in the apparent plaid speed of 5.66°/s (marked by empty arrows). To assess the effect of stimulus speed on transfer of learning, we also included a grating whose speed matched the plaid speed of 5.66°/s (Be, empty arrows). Although all stimulus conditions were conducted together, we analyze and present data into two batches: bidirectional transfer between component and pattern motion (Figure 5) and transfer to other stimulus features (Figure 6).

For the plaid-training group, the plaid stimuli consisted of two orthogonal component gratings (component contrast = 50%, size = 8°, 2D raised cosine spatial envelope; component spatial frequency = 1 cycle/°; Figure 3Ba). Component drift speed was 4°/s, which resulted in the plaid velocity of 5.66°/s. Training motion directions were either left/right or up/down, counterbalanced across participants. In addition to the trained condition, duration thresholds were measured for five additional pre- and post-test stimulus conditions: (1) a moving grating with the same apparent direction and speed as the trained plaid stimulus (Figure 3Bb); (2, 3) two component gratings that constituted the trained plaid stimulus (i.e., gratings with direction ±45° away from the trained directions; Figure 3Bc. Note that these data were collected in two separate blocks, each testing one motion direction axis, and subsequently averaged to get a single threshold estimate); (4) a plaid stimulus moving to the untrained directions, but comprised of same static component features (Figure 3Bd); (5) a grating moving in the trained directions (left/right) but with the original plaid apparent speed (speed = 5.66°/s, Figure 3Be).

### Experimental procedures and Data analysis

Pre- and post-test consisted of six randomly ordered blocks corresponding to different stimulus conditions (the trained stimulus, plus 5 additional stimulus conditions, as described above). In each block, stimulus durations were controlled by two 80-trial interleaved staircases (a 2-down-1-up staircase and a 3-down-1-up staircase), yielding 160 trials for each threshold estimate. The initial starting durations for two staircases were 100 ms and 110 ms, respectively. Pre- and post-test measurements were conducted on day 2 and day 7, respectively. On day 1, each participant completed a practice phase that was identical to the pre- and post-test battery, except that each block consisted of only 60 trials. The purpose of this practice day was to help stabilize pre-test measurements. The perceptual training lasted four days (days 3-6). On each day, participants completed seven 100-trial blocks, resulting in a total of 28 training blocks. For the first training block on the first training day (day 3), the initial starting durations for the two staircases were 100 ms and 110 ms. For all subsequent training blocks, the initial stimulus durations were the durations in the final trials of two staircases in the previous training block. All participants completed these seven experimental sessions within 14 days.

To estimate duration thresholds for each pre- and post-test condition, we fit Weibull psychometric functions to 160 trials of raw data using the maximum likelihood method, estimating the thresholds at 82% correct. The amount of learning in each condition was estimated by computing percent of improvement (PI):

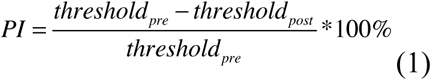

where *threshold*_*pre*_ and *threshold*_*post*_ indicate duration thresholds for the corresponding pre- and post-test stimulus conditions. We used paired t-tests for comparisons of pre- and post-test thresholds and for comparison of PI across stimulus conditions. One-sample ttests were used for assessing the statistical significance of PI against the null hypothesis of 0% PI. All t-tests were two-tailed and performed using Matlab Statistical and Machine Learning Toolbox.

**Figure 4.**
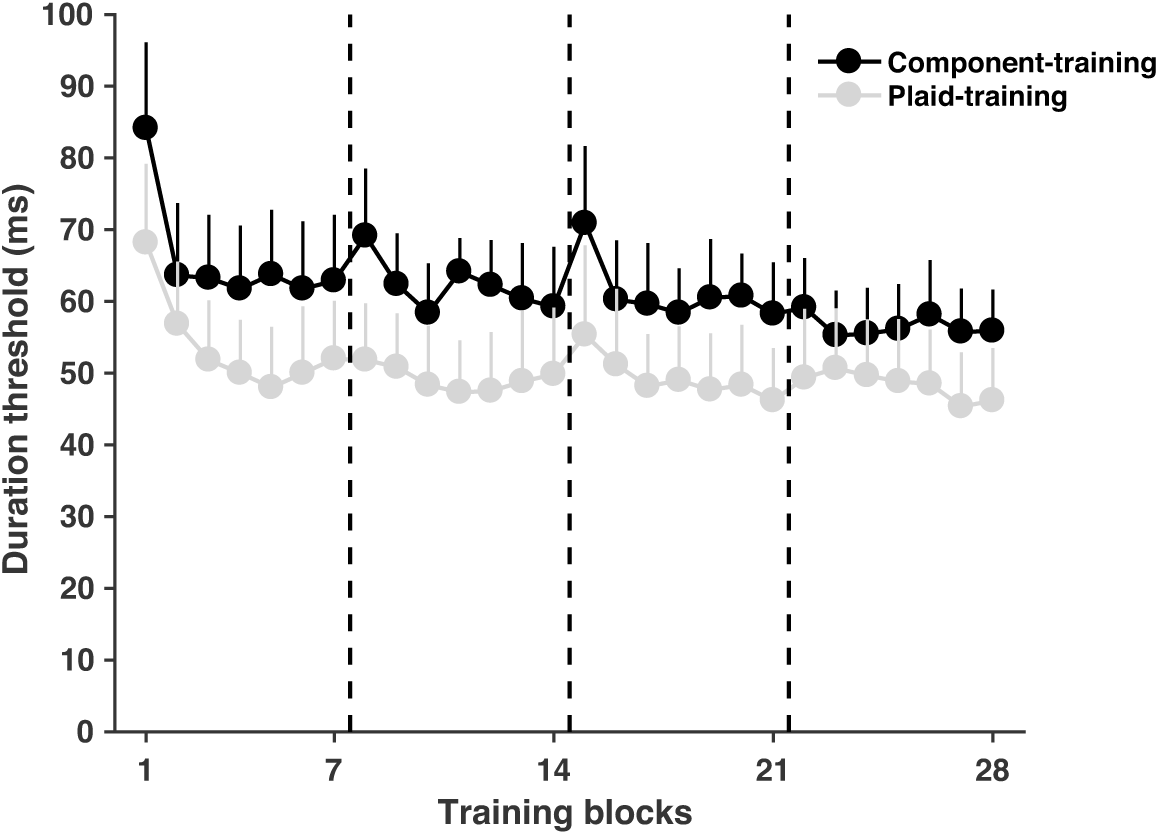
Learning curves for the component-(black) and the plaid-training (gray) group. Data are thresholds for 28 training blocks, tested over 4 days of training. Vertical dashed lines separate data for four training days. Note that the plaid-training group showed lower duration thresholds. This is expected given the faster apparent speed of plaid stimuli and known effects of stimulus speed on temporal duration thresholds ^36^. Error bars are SEM across subjects.

## RESULTS

### Effective perceptual learning for both component and plaid stimuli

We first examined whether our training procedure was sufficient to result in perceptual improvement. Here, for each group, we compared pre- and post-test thresholds for the trained stimulus condition. The results revealed significant improvements in thresholds for both the component- and the plaid-training group (Figure 5E; t(7) = 2.79, p = 0.0268 and (5) = 6.28, p = 0.0015, respectively). We also computed percent of improvement (PI, see Equation 1), and found significantly positive PIs for both groups (Figure 5F; t(7) = 5.06, p = 0.0015; t(5) = 12.04, p = 6.97 x 10^−6^), with each group showing about a 20% improvement in performance.

**Figure 5.**
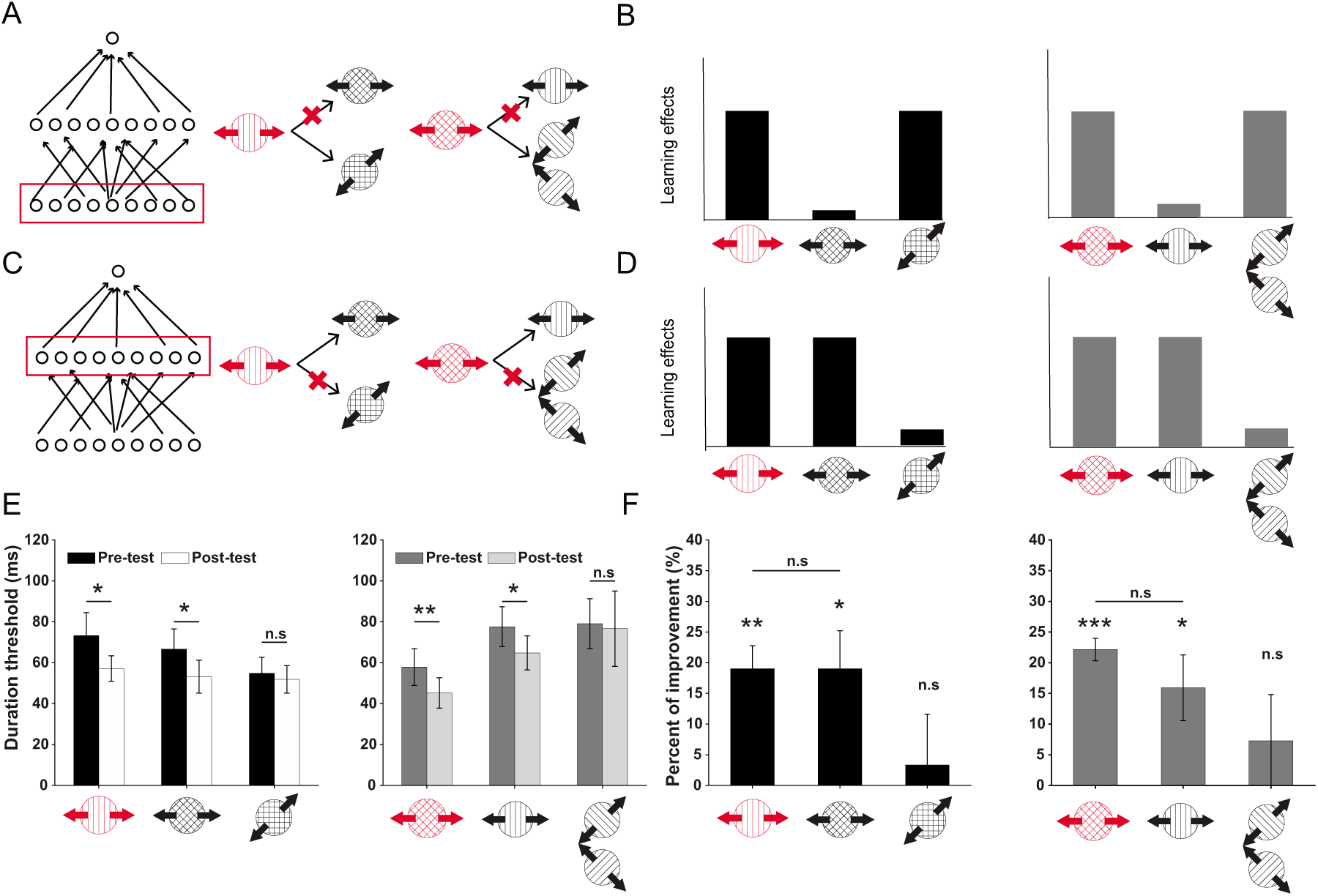
Schematic illustrations (A, C), predictions (B, D) and empirical results (E-F) for component-dependent and pattern-dependent VPL. ***A***. The component-dependent VPL takes place at the lowest level of motion processing, as indicated by the red rectangle. Here, training on a component stimulus should only transfer to the plaid stimulus that comprises the trained component. Moreover, training on a plaid stimulus should only transfer to its two constituent components. ***B***. Learning effects as predicted by component-dependent learning in panel A. ***C-D.*** Illustrations of the pattern-dependent perceptual learning and its predicted learning effects, following conventions in panels A and B. Here, plasticity takes place at the middle stage of motion processing. (E) Duration thresholds at pre-/post-test across stimulus conditions in the component (left panel) and the plaid training (right panel), respectively. (F) Learning effects quantified as percent of improvement (PI%) across stimulus conditions and training regimes. The overall pattern mimics the predictions in (D), indicating that plasticity likely occurs at the middlelevel of motion analysis. For all subplots, error bars denote ±1 SEM across subjects. Significance symbol conventions are *:p < 0.05; **:p < 0.01; ***: p < 0.001; n.s.: non-significant. Same definitions of error bars and symbol conventions are kept for all figures in this paper.

### Bidirectional transfer of learning between component and plaid motions

The main focus of this paper is to examine the transfer of perceptual learning to a range of diagnostic stimulus conditions. A two-stage criterion was used to assess transfer of learning. First, we concluded that learning transfers to a stimulus condition if the pre/post-test difference on this condition was statistically significant. If a stimulus condition passed this first test, then we compared its PI to the corresponding trained condition (i.e., either trained component or trained plaid). If the transfer PI was significantly smaller than the trained PI, the result was described as a “partial transfer”. Alternatively, if the PI for a transfer condition was not statistically smaller than the PI for its corresponding trained condition, we referred to it as “complete transfer”, according to an established convention in VPL research ^13,16,17^.

The key aim of this study was to determine whether perceptual training leads to plasticity within low-level component-dependent motion processing or middle-level pattern-dependent motion processing. To be precise, component-dependent plasticity predicts that training on a component motion stimulus should only transfer to the plaid composed of the trained component gratings, and that training on a plaid stimulus should only transfer to its two constituent components (Figure 5A-B). On the contrary, pattern dependent plasticity predicts that training on a component motion stimulus or on a plaid motion stimulus that moves in the same directions should mutually transfer to each other (Figure 5C-D).

Our results were consistent with plasticity in pattern-dependent mechanisms. First, perceptual training on a component grating significantly reduced the duration thresholds on the plaid that moved in the same apparent direction as the trained grating (Figure 5E left panel, pre-/post-test, t(7) = 2.88, p = 0.0237; Figure 5F left panel, PI, t(7) = 3.08, p = 0.0178). More importantly, the PI was statistically equivalent to the PI on the trained grating (Figure 5F left panel; t(7) = 0.002, p = 0.999). Consistently, perceptual training on a plaid stimulus also transferred to the component grating that moved in the same apparent direction (Figure 5E right panel, pre-/post-test, t(5) = 3.336, p = 0.0207; Figure 5F right panel, PI, t(5) = 2.971, p = 0.0311). Also, the PI on the untrained component was not statistically different from the PT on the original trained plaid (Figure 5F right panel, PI, t (5) = 1.29, p = 0.2533). The bidirectional transfer between the component and the plaid stimuli that moved to the same directions suggest that perceptual training most likely alters the computation in the visual units that process the pattern motion direction. Moreover, training effects on a component did not significantly transfer to a plaid that included the trained component, but moved in a different direction (Figure 5E left panel, pre-/post-test, t (7) = 0.784, p = 0.4586; Figure 5F left panel, PI, t(7) = 0.405, p = 0.6978). Plaid training also did not improve the performance on its two constituent components (Figure 5E right panel, pre-/post-test, t (5) = 0.305, p = 0.7709; Figure 5F right panel, PI, t(5) = 0.963, p = 0.3797). Taken together, these findings suggest that pattern-dependent learning at the middle-level motion system, rather than component-dependent learning at the low-level motion system, plays a pivotal role in mediating learning transfer of motion.

### Specificities to direction, speed, size, and contrast

We have thus far focused on experimentally disentangling component-dependent from pattern-dependent VPL, with the results arguing against low-level component-dependent VPL. What remains unclear, however, is whether the perceptual training led to enhancements in the processing of sensory features or high-level non-sensory attributes. For instance, participants might learn motion directions as abstract concepts ^16^ or be more familiar with the general task statistics (e.g., stimulus timing, stimulus-response association ^17^). In this case, plasticity takes place in higher brain hierarchy that is independent of the sensory processing. To further delineate the plasticity in the sensory (Figure 6A-B) or the non-sensory processing (Figure 6C-D), we examined the tolerance of our training across several other forms of stimulus variations, i.e., direction, speed, size, and contrast. The prediction is that if the plasticity is largely limited to sensory processing, learning should be confined to the trained stimuli; otherwise learning effects will transfer irrespective of the variations in other stimulus features.

**Figure 6.**
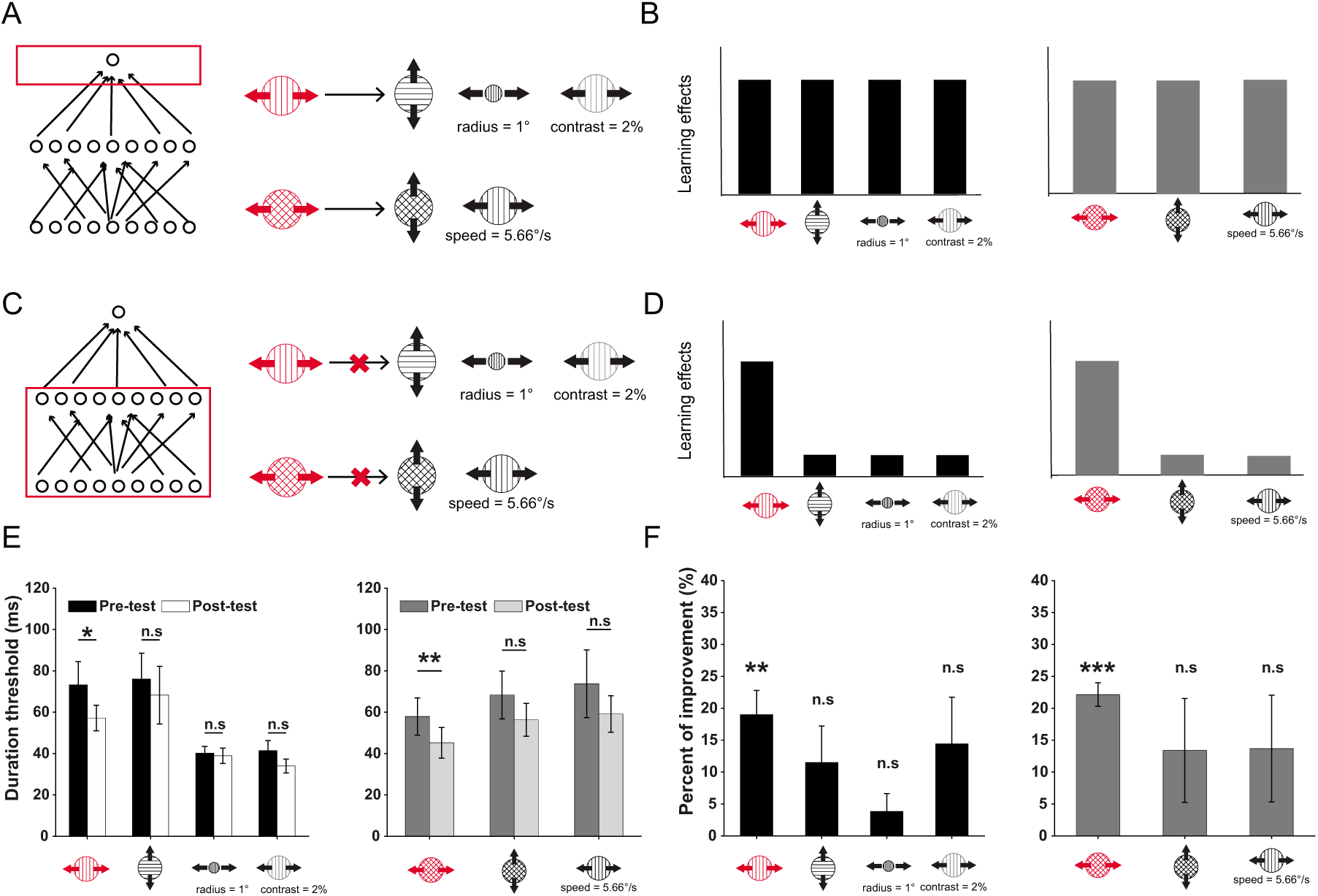
Specificity of motion VPL to basic sensory features. ***A-B***. Illustrations and predictions of the plasticity at the highest-level stage (e.g., PFC) in the three-layer network. This mechanism predicts that training on a component or a plaid stimulus should be generalizable regardless of the variations in low-level visual features, such as direction, speed, size, and contrast. ***C-D*** Illustrations and predictions akin to (A-B), expect that the plasticity occurs within the general sensory representation stage. This scheme predicts that training on a component or a plaid stimulus should exhibit minimal transfer to the stimuli that differ in basic visual features. (E) Duration thresholds at pre-/post-test across stimulus conditions in two training groups. (F) Empirical learning effects, quantified as percent of improvement (PI%), across stimuli and training groups. The transfer pattern of learning is more consistent with predictions in (D). No significant transfer in all other stimulus conditions is noted, implying the plasticity within the sensory representation level as shown in (C).

The results indicated a notable specificity to stimulus variations. In the component-training group, we did not find significant transfer for trained and test stimuli that differed in motion directions (Figure 6E left panel, pre-/post-test, t(7) = 1.886, p = 0.101; Figure 6F left panel, PI, t(7) = 2.016, p = 0.084). We also found no significant transfer to test stimuli that have smaller size (Figure 6E left panel, pre-/post-test, t(7) = 1.308, p = 0.232; Figure 6F left panel, PI, t(7) = 1.376, p = 0.211) or lower contrast (Figure 6E left panel, pre-/post-test, t(7) = 2.187, p = 0.065; Figure 6F left panel, PI, t(7) = 1.971, p = 0.089).

Similarly, if component motion directions were switched such that the resulting plaid moves in an orthogonal direction, transfer effects in the plaid-training group were not statistically evident (Figure 6E right panel, pre-/post-test, t(5) = 1.268, p = 0.261; Figure 6F right panel, PI, t(5) = 1.645, p = 0.161). We also investigated how changing stimulus speed affects learning transfer. When the grating speed was increased to match the apparent speed of the trained plaid, the transfer effect was not significant (Figure 6E right panel, pre-/post-test, t(5) = 1.257, p = 0.265; Figure 6F right panel, PI, t(5) = 1.635, p = 0.163).

Taken together, we find that motion VPL is specific to stimulus direction, speed, size, and contrast. These results demonstrate that our training has strong susceptibilities to variations in basic visual features. Such strong dependencies indicate that a broadly tuned non-sensory learning mechanism unlikely plays an important role in observed learning because it predicts a broad transfer over variations in low-level stimulus features. Note that we cannot completely eliminate the possibility of changes in sensory readout mechanisms since, theoretically, a refined readout mechanism can be sensitive to changes in sensory features ^37,38^. Nonetheless, these results suggest the pivotal roles of basic stimulus features in perceptual learning of motion.

## DISCUSSION

Elucidating where in the visual processing hierarchy plasticity associated with VPL takes place has been a key question in perceptual learning research over the past decades. Here, we addressed this question in the domain of motion perception. We trained participants to identify motion directions of either component motion (a drifting grating) or pattern motion (a drifting plaid), and assessed transfer of learning to a variety of carefully controlled stimulus conditions. The bidirectional transfer of learning between component and pattern motion provides evidence that learning effects most likely take place at the middle-levels of processing where component motions are combined into plaid percepts, and, at the same time, rules out plasticity at the low-levels where complex motions are represented as components. In addition, we also observed specificities to the trained direction, speed, stimulus size, and contrast. These results are in line with the previous findings that VPL is generally vulnerable to the variations in basic feature dimensions and argue against plasticity in high-level brain areas that represent non-sensory cognitive factors, such as general task statistics and decision rules ^15,16,17^.

Our results are of significance for understanding mechanisms underlying motion perception. As one of the key research topics in vision science, dissociable functional roles of the low-level and the middle-level motion system have been well documented ^22,39,40^. The seminal paper by Adelson and Movshon ^32^ documented how moving plaid percepts can arise from component gratings. Subsequent neurophysiological work discovered distinct tuning properties of individual neurons in V1 and MT with preferences toward component and plaid representations, respectively ^30^. These findings were generalized to humans. Huk and Heeger ^41^ reported robust fMRI adaptation to pattern motion in the human motion-sensitive area hMT+. Thus, the phenomenon of component and pattern motion serves as a good benchmark for studying visual hierarchy of motion processing.

Although we have a good understanding of visual motion processing hierarchy, we know little about the roles different stages play in VPL. We address this question by showing that training on component or pattern motion bi-directionally transfers to each other if the two stimuli share the same apparent motion direction. These results suggest that, when a plaid motion stimulus is being learned, learning signals might preferentially refine the pattern-selective units that respond to the apparent motion direction, but not the component sensitive units. While there have been many behavioral studies of motion VPL, to our knowledge, no studies employed an experimental design that allowed distinguishing between plasticity at low and at the middle levels of motion processing. For instance, VPL studies typically relied on random-dot-kinematogram stimuli or trained subjects on fine direction discrimination tasks ^7,42,43^. Studies that used gratings only tested contrast thresholds for coarse motion direction judgments ^44^.

Our study also constrains theoretical models of VPL. Two distinct computational frameworks of VPL have emerged so far, where learning either improves the quality of sensory encoding or optimizes high-level readout and decision mechanism that can in turn promote perceptual sensitivity. Empirical evidence, however, is highly contentious. Early psychophysical studies on motion VPL demonstrated the considerable specificity to the trained direction ^7,8^, implying the plasticity among direction-selective units. However, specificities in motion VPL have also been shown to be mediated by other factors, such as task difficulty ^45,46^, exposure to other directions ^47^, external noise ^44^. This debate in VPL psychophysics is mirrored by a similar debate with respect to the neural substrates of VPL. For example, after training monkeys on a motion direction decision task, Law and Gold ^27^ found pronounced behaviorally relevant changes in neural responses in area LIP, but minimal changes in neural activities in area MT. This study advocates a mechanism beyond the sensory-representation level, where training results in a more efficient extraction of useful sensory information rather than in an enhancement of sensory representations per se. In contrast, recent fMRI studies found that motion VPL refines the cortical tuning of the human MT, emphasizing the pivotal role of enhancement at sensory-representation level ^48,49^. Notably, the mechanistic role of high-level cognitive influences in sensory processing is still largely unknown. Previous studies have suggested at least two broad categories, mechanisms that are sensory (e.g., selective readout) and those that are non-sensory (e.g., conceptual learning, rule-based learning). While disentangling between these higher level processes is beyond the scope of this paper, the observed specificity to basic stimulus features argues against non-sensory cognitive factors.

What are the possible neural underpinnings of the observed empirical findings in the present work? We surmise that several mechanisms may coexist and interact. First, because training on a plaid motion stimulus does not fully transfer to its two components (Figure 5E), we conclude that a significant part of the relevant plasticity occurs downstream from the low-level motion mechanisms. Given the evidence that MT neurons analyze pattern motion by selectively integrating inputs from a population of V1 neurons ^38^, one possible mechanism is that learning improves information transmission from the low-level to the middle-level motion processing. Such a mechanism is consistent with findings of a recent study where attention was shown to improve the amount of information transferred from V1 to hMT+ ^50^. Moreover, learning effects in our study are specific to direction, speed, contrast, and size, indicating critical roles of neuronal tuning to these low-level visual features. For example, stimulus contrast and size have strong influences on neural responses in motion processing ^51^. This is also in line with our previous findings showing that motion perception is strongly modulated by stimulus contrast and size ^52,53^—behavioral findings that have been linked to mechanisms within area MT ^54,55^.

In summary, our study provides evidence for the training-induced plasticity in the intermediate stage of motion processing, and highlights the significance of basic motion-related visual attributes in mediating the transfer of motion VPL.

